# Genomic Environments and Their Influence on Transposable Element Communities

**DOI:** 10.1101/667121

**Authors:** Brent Saylor, Stefan C. Kremer, T. Ryan Gregory, Karl Cottenie

## Abstract

**Background:** Despite decades of research the factors that cause differences in transposable element (TE) distribution and abundance within and between genomes are still unclear. Transposon Ecology is a new field of TE research that promises to aid our understanding of this often-large part of the genome by treating TEs as species within their genomic environment, allowing the use of methods from ecology on genomic TE data. Community ecology methods are particularly well suited for application to TEs as they commonly ask questions about how diversity and abundance of a community of species is determined by the local environment of that community.

**Results:** Using a redundancy analysis, we found that ~ 50% of the TEs within a diverse set of genomes are distributed in a predictable pattern along the chromosome, and the specific TE superfamilies that show these patterns are relate to the phylogeny of the host taxa. In a more focused analysis, we found that ~60% of the variation in the TE community within the human genome is explained by its location along the chromosome, and of that variation two thirds (~40% total) was explained by the 3D location of that TE community within the genome (i.e. what other strands of DNA physically close in the nucleus). Of the variation explained by 3D location half (20% total) was explained by the type of regulatory environment (sub compartment) that TE community was located in. Using an analysis to find indicator species, we found that some TEs could be used as predictors of the environment (sub compartment type) in which they were found; however, this relationship did not hold across different chromosomes.

**Conclusions:** These analyses demonstrated that TEs are non-randomly distributed across many diverse genomes and were able to identify the specific TE superfamilies that were non-randomly distributed in each genome. Furthermore, going beyond the one-dimensional representation of the genome as a linear sequence was important to understand TE patterns within the genome. Additionally, we extended the utility of traditional community ecology methods in analyzing patterns of TE diversity.

## Background

### Transposable Elements in the Genome

Transposable elements (TEs) are mobile genetic elements that comprise a large portion of most eukaryotic genomes. The human genome, for example, contains more than 3 million copies of various types of TEs, making up between half and two thirds of the total quantity of DNA [1]. The diversity and abundance of TEs in the genome is influenced by coevolution with the host, and the interaction between properties of the genome and properties of the TEs. For example, some TEs persist because they have been co-opted for important regulatory or structural functions [2–4], whereas others are known as disease-causing mutagens [5–7] that remain abundant as a result of their ability to make copies of themselves, despite their detrimental effects on the host genome [8,9]. In this regard, the relationship between TEs and their host genomes may be considered along an ecological continuum from mutualism at one extreme, through commensalism, to strict parasitism at the other end of the spectrum [10].

Beneficial TE insertions can be preserved by natural selection acting at the host level [11,12], and others may accumulate via mutation pressure (e.g., if net insertions outweigh TE deletions) or genetic drift (especially in small populations) if they simply do not exert a significant negative impact on host fitness [13–15]. However, TEs that impose fitness costs on the host, either as deleterious mutagens or simply as extra genetic baggage, can persist in the long term only if they are able to evade inactivation by the host. This can be done by a combination of replicating more rapidly than they are silenced by host defenses [16–18] and/or spreading more quickly than they are removed from the population via host-level purifying selection [19,20].

These process result in substantial variability in the diversity and abundance of TEs within and among genomes. Within individual genomes, TE diversity can be seen in the large number of TEs and TE superfamilies (1355 and 37 respectively in humans [21]). The variation in TEs between genomes is even more evident, with the number of TE superfamilies ranging from 1 in *Dirofilaria immitis* (dog heartworm) to 39 in *Branchiostoma floridae* (lancelet), *Bombyx mori* (silkworm moth), and *Hydra magnipapillata* (freshwater hydra) [22], and abundance of individual TEs varying widely, even among individuals of the same species [23,24].

### Types of Transposable Elements

In addition to varying in abundance and distribution, TEs are mobile within the genome, and are grouped into two broad classes based on how the move/transpose. Class I TEs, or retrotransposons, move within the genome using a copy-and-paste mechanism. Copy-and-paste transposition involves transcription of TE DNA into an RNA intermediate, with the element itself remaining in its original location and serving as a template. The RNA intermediate then exits the nucleus where it is reverse transcribed into DNA, which then re-enters the nucleus and inserts into a new location [25–28]. Most Class II elements transpose using a cut-and-paste mechanism without an RNA intermediate. Cut-and-paste transposition involves excising the TE itself from its location in the genome for reinsertion into a new location without leaving the nucleus. Increased copy number in cut-and-paste transposons is accomplished by the repair mechanisms of the host genome responding to the breaks left behind by the TEs excisions, which fills in the missing DNA from the complementary strand. Heletron and Maverick elements are Class II elements that use a form of cut-and-paste transposition with no RNA intermediate [29].

### Evolution, Ecology, and the TE Perspective

These factors – TE effects on host fitness, suppression by the host genome, TE accumulation and dispersal within (and between, see for instance horizontal migration [30]) genomes, and the evolutionary relationships between the different TE families [31] – are all important in shaping TE abundances in different genomes. Some of these factors, such as the phylogenetic relationships between the TE families and the coevolution between TEs and host mechanisms for suppressing TE replication, are explicitly evolutionary from the perspective of the TE. Other processes, such as the dispersal of TEs to other parts of the genome, are explicitly ecological from the perspective of the TE. According to Linquist *et al.* [32], an explanation is ecological if it focuses on changes in composition of the community of TEs and how TEs interact with the host genome and other elements in it, whereas evolutionary explanations relate to changes in the TE sequences themselves over generations, including co-evolution with the host genome. This distinction becomes important when the mechanisms responsible for shaping TE distribution or abundance can be either evolutionary or ecological. For instance, an ecological explanation for the accumulation of TEs in a specific area of the genome could be that that area is available and the nearby TEs are able to disperse there. However, if a specific group of TEs changed in a way that let them exploit that same area, that would be an evolutionary explanation for that same observation.

The notion of “genome ecology” has been invoked numerous times in the TE literature, however, many of the purported examples actually relate to TE *evolution*, and conflating TE ecology and TE evolution can result in asking the wrong questions and using the wrong tools [32]. In a recent study, we applied an explicitly ecological approach to the analysis of patterns of TE distribution and abundance within the genome of the cow, *Bos taurus* [33]. Specifically, TE distribution was assessed using a well-established method derived from community ecology, akin to assessing community composition along an environmental gradient (e.g., how communities might vary along a mountain range; see e.g., Whittaker [34]). Our genomic version of this analysis examined how communities of TE superfamilies varied along each chromosome.

To implement this community ecology approach in the study of TE distribution, each chromosome in the cow genome was divided evenly into discrete windows. The abundance and diversity of the TE superfamilies in each window was then assessed. Combining the superfamily counts in each window resulted in a TE community for each window. Various properties of the genome could then be examined as potential correlates of variation in local TE community composition. In Saylor *et al.*[33], we considered the location of the window along the chromosome and local gene density as predictors, and these were used to test if the TE communities of each chromosomes changed in predictable ways from one window to the next. The results in *Bos taurus* found that 50% of the within-chromosome variation in TE community composition was explained by examining physical position along that chromosome [33]. Our analysis demonstrated the power of this ecological approach, but it was largely a proof-of-concept and examined only one genome. Moreover, we implemented the most straightforward way to measure the location of any genetic element: the location of their sequence along the chromosome.

### TEs in a 3D Environment

The above approach is the most intuitive way to represent loci on a chromosome. Most chromosome maps (physical, genetic, and karyotype) are linear in nature; however, this is an oversimplification of the actual distance between two loci on a linear chromosome. These idealized maps are representative of the phases of the cell cycle associated with replication, which make up a relatively short part of the cell’s life cycle [35]. During interphase, which makes up the majority of the cell’s lifecycle, chromosomes are found uncondensed within the nucleus, where they are arranged into chromosome territories (CT) [36–39]. Each CT contains one chromosome, with interaction between chromosomes occurring at the borders between territories. CTs can be further divided into genomic compartments and subcompartments [40]. Genomic compartments are made up of alternating segments of heterochromatin (tightly-bound, less accessible DNA) and euchromatin (loosely bound, more accessible DNA) [39]. Genomic subcompartments are areas within a genomic compartment that physically interact more often with each other than one would expect by chance. Although the specific reasons these subcompartments form is not clear, each of the six subcompatments identified by Rao *et al.* (2014) has a distinct histone modification pattern and interaction profile, indicating that they are regulated in similar ways.

This physical 3D structure of a chromosome within a cell has important implications for how genes are expressed, how they interact with regulatory elements, and how accessible the DNA is to proteins [41–43]. Like genes, TEs also need to be accessed by regulatory elements, transposases, and polymerases to function. It is thus likely that it will also influence the TE distribution along the chromosome. This, however, has never been studied. If physical structure does have an influence on TE community dynamics, we would expect sub-compartments that are physically close to each other will be more similar than predicted based on chromosome location alone. This is consistent with findings by Sanyal *et al.* [44], who found only 7% of looping interactions are with the closest gene, and a strong correlation between long range promoter-enhancer interaction and gene expression. If genomic subcompartments are acting as different genomic environments, we would also expect heterochromatic regions (subcompartments B1-B4) and euchromatic regions (subcompartments A1-A2) to have different TE community compositions. If these chromosome structural properties are important determinants of TE location, we would expect strong relationships between these properties and TE locations along the chromosome, similarly to the analysis of the *B.taurus* genome [33].

While this relationship between functional chromosome structure and TE chromosomal distribution is the primary focus of this study, we will also study the generality of these TE spatial patterns. Chromosome subcompartment data were only available for the human genome, as it is the only chromosome interaction analysis with a sufficiently high resolution to detect subcompartments [40]. To confirm our results in other genomes, we will also explore the generality of spatial findings across a suite of species with genomes with high sequencing depth, scaffolds assembled into full chromosomes, and well-annotated TEs. There are 11 species that fit these criteria available from Genbank.

Chromosome structure within genomes appears to be a universal genome property, from chromosome territories at the coarsest level of organization [45], to chromosome looping which has been found in a wide variety of prokaryotes and eukaryotes [46]. If the TE spatial distribution is (partially) determined by these universal chromosome structural properties, then we predict that the spatial patterns in TE distributions would be detected across a wide variety of genomes.

To replicate the spatial analysis across multiple species, one methodological issue must be solved first. In the primary analysis of Saylor *et al.* [33], the window sizes were determined independently for each chromosome so that each window contained an average of 100 TEs. This ensured that a TE community would be examined within each window, no matter the size of the chromosome. However, it had the less-desirable effect of normalizing TE density as a chromosome property, possibly obscuring TE density itself as an explanatory factor of the importance of spatial location. Using a systematic approach with evenly distributed fixed window sizes should make it possible to identify the effect of window size on this analysis. The Saylor *et al.* [33] study used a fixed window size across the entire genome. The fixed window method produced very similar results to the dynamic window approach when the fixed window size was similar to the average size of the dynamic windows, with the added benefit that windows on any chromosome, in any genome, were directly comparable. However, since the window size affects the number of communities on each chromosome, the average size of each individual community of TEs, and in turn the computational resources required to conduct the analysis, were very different. How to choose an appropriate window size for the analysis of a given genome was not fully explored. Additionally, it remains to be determined whether similar window sizes can be used across widely different genomes in such a way that the results can be compared.

The present study investigates the importance of a chromosome’s 3D spatial structure (the “genomic environment”) on the distribution of the TEs on each of the chromosomes in the human genome and assesses the usefulness of using TEs as indicators of specific genomic environments. Additionally, we investigate 11 genomes from diverse eukaryotic organisms to investigate if variation in the TE community can be explained by where it is in the genome is a general property of TE communities. To accomplish this, we also assess the impact of window size on the detecting the spatial structure of the TE community within each of our 11 study genomes.

## Methods

### Study Species and Genome Data

Of the available whole genome sequences, only those that were assembled into chromosomes were considered. Eleven eukaryote genome species – including representative vertebrates, invertebrates, plants, and fungi – were included in the present study, on the basis of genome size, chromosome number, and phylogenetic diversity. These are summarized in Table 0.1. Where available, the reference assembly was used. If not, the representative assembly was used where possible. If neither one of those was available, the most recent assembly was used. The output of RepeatMasker searches of each genome were downloaded from the Genbank FTP site. These files contain the names and locations of any region in each genome that matched TEs in the RepBase TE database.

**Table 0.1.** Summary of the 11 species included in the present analysis, including information on genome size, chromosome number, and source sequence accession.

| *Species* | *Common name* | *Genome size(bp)* | *Chromosome number (n)* | *Assembly accession* |
| --- | --- | --- | --- | --- |
| *Homo sapiens* | Human | 3088269832 | 23 | GCF_000001405.28 |
| *Bos taurus* | Cow | 2670123310 | 30 | GCF_000003055.6 |
| *Mus musculus* | Mouse | 2730855475 | 21 | GCF_000001635.23 |
| *Drosophila melanogaster* | Vinegar fly | 143706478 | 7 | GCF_000001215.4 |
| *Takifugu rubripes* | Puffer fish | 391484725 | 22 | GCF_000180615.1 |
| *Danio rerio* | Zebra fish | 1371702787 | 25 | GCF_000002035.5 |
| *Anopheles gambiae* | Mosquito | 265011681 | 5 | GCF_000005575.2 |
| *Caenorhabditis elegans* | Roundworm | 100272607 | 6 | GCF_000002985.6 |
| *Arabidopsis thaliana* | Mustard weed | 11914634 | 6 | GCF_000001735.3 |
| *Oryza sativa* | Rice | 382150945 | 12 | GCF_000005425.2 |
| *Zea mays* | Maize | 2059701728 | 20 | GCF_000005005.1 |

### Transposable Element Categorization

The bins used to categorize the TEs within each genome are based on the output from the Genbank run of RepeatMasker on each genome. Our analysis used the superfamily level classification of TEs as it is the most well-defined classification below the more general TE Class. It is also the most commonly reported, which increases the degree to which data can be compared across these different genomes. Modifications to the RepBase classification were carried out to reflect updates to TE superfamilies subsequent to the record submission to Repbase [47–49]. Several groups of LINEs, SINEs, LTR retrotransposons, and DNA transposons could still not be identified to the superfamily level in their original publications. These will be referred to as superfamilies for simplicity, however, they reflect less well-defined groupings.

### Quantifying Within-Chromosome Spatial Community Structure

To detect the relative impact of within-chromosome community structure for all of the spatial analyses, we used redundancy analysis (RDA) as implemented by the vegan package in R [50]. This performs a multivariate multiple regression with the counts for the number of TEs in each window as the dependent variable, and the properties of the genomic/chromosomal environment as the independent variable. This results in an R^2^ value for each chromosome which represents how well the TE community in each window can be predicted based on the environmental variable used. The spatial environmental was modelled with the principal components of neighbouring matrices (PCNM) [51,52] procedure. The input for the PCNM for the analyses that use linear spatial structure was a dissimilarity matrix representing the distances between each window, and the input for the 3D analysis was a dissimilarity matrix based on the interaction frequencies from the HiC data of each chromosome. For more details, see Saylor *et al.* [33].

### Window Size Analysis

This analysis used RDAs of TE abundances across windows as a function of spatial location of the window to explain TE communities within each chromosome as above and in Saylor *et al.* [33]. This was done for each chromosome in each of the 11 genomes at each of 20 different window sizes ranging between 10x the size of the largest *Bos taurus* TE (14,753bp) at minimum to the size of the smallest *Bos taurus* chromosome at a maximum (4,404,134bp). This resulted in windows ranging from 14,753bp to 4,404,134bp by increments of 219,469bp.

### Genomic and Chromosomal Properties

In in addition to assessing the impact of window size on the detection of spatial patterns in TE community composition along the chromosome, we also assessed the impact of changing the window size on chromosomes with different environmental properties. we selected one “large” (790,718bp) and one “small” (144,808bp) window size because they represented extremes of window sizes while avoiding sizes small enough to cause statistical issues (see Discussion). The genomic properties investigated with these two window sizes were: 1) The total length of the available genome sequence; 2) The C-value, an independent genome size estimate of physical size for that species taken from the Animal Genome Size Database [53] or Plant DNA C-values Database [54]; 3) the difference between the genome size estimate and the available sequence length, which serves as a measure of how complete the sequence is; and 4) the number of chromosomes. The chromosomal properties, downloaded from the Genbank entries for each genome (Supplementary table 1), were: 1) Genome, which consisted of which genome the chromosome was from and was used in the phylogenetic independent contrast to account for non-independence of the data; 2) Chromosome length, which consisted of the length of the sequence for that chromosome and was used to measure the amount of space for the TEs to insert; 3) GC%, the percentage of the sequence made of guanine-cytosine basepairs, which is highly variable, easily calculated from the sequence, and has been correlated with the presence of some TE families and other genomic features (see Eyre-walker & Hurst, 2001 for review); 4) Number of genes, which directly measures the number of genes on the chromosome; and 5) Number of proteins, which measures the number of those genes with known or putative protein products. Each of the genomic properties was compared to average R^2^_adj_ for all of the chromosomes in that genome and each of the chromosomal parameters were compared to the R^2^_adj_ for each chromosome across genomes.

The phylogenetic distances between the 11 host species were downloaded from the sequenced tree of life webpage [56]. The resulting phylogeny was used to run a phylogenetically independent contrast (PIC) analysis on the R^2^_adj_ from the RDA for each chromosome, and on the genomic and chromosomal properties. In the chromosome property analysis, polytomies were added to the tips with each chromosome in each genome being equally related to each other. The contrasts of the average R^2^_adj_ values were then compared to the contrasts of the genomic properties as above.

### TEs in a 3D Environment

10kb resolution interaction frequency data with MAPQ scores above 30 generated by Rao *et al.* (2014) were downloaded from the Gene Expression Omnibus (GEO) database (GEO accession GSE63525). These frequency data were KR normalized to adjust for methodological artifacts according to the instructions downloaded with the data. The genomic compartment locations were calculated by taking the first principal component of the normalized interaction matrix [57], using the cmdscale function in the stats package of R.

The genomic subcompartment data were also downloaded from the Rao *et al.* dataset hosted in the GEO database. The subcompartment data consisted of start and end positions for each subcompartment along the sequence of each human chromosome, and which of the 7 subcompartments types (A1, … NA) that section is classified as. We then associated that structural information to our TE distribution data. For each window in our chromosome spatial analysis, we calculated the proportion of the window made up of each subcompartment. This resulted in 7 variables consisting of the proportion for each window made up of that subcompartments.

Finally, we repeated the spatial RDA for the human chromosomes (see Quantifying within-chromosome spatial community structure section above), but this time in addition to the spatial patterns obtained with PCNMs, we used spatial patterns generated by PCNMs of the frequency data, the 7 additional explanatory variables from the subcompartments, the first principal component of the interaction frequency matrix, and the number of genes in each window.

### Indicator Species Analysis

The usefulness of each TE, and each pair of TEs as an indicator of genomic subcompartment in the human genome was determined using the multipatt function found in the indicspecies package for R [58–60]. This analysis computes two types of an indicator entity: Indval, which evaluates the strength of using each TE as an indicator of a specific environment; and Phi, which is a measure of correlation between the species presence/absence matrix, and the genomic subcompartment. Each of these statistics was measured for each TE in each genome.

IndVal scores range from 0 to 1, with 1 indicating a TE always occurs in a given environment, and never occurs in other environments, and 0 indicating a given TE never occurs in a given environment and is always found in other environments. Within each chromosome IndVal scores were generated for each TE for each subcompartment / pair of subcompartments. The significance of each score was assessed using a permutation test, and significant scores, where *p* < 0.05 were reported.

Phi scores were also produced for each of the 22 human chromosomes. Phi scores also range from 0 to 1, with 1 being perfect correlation between two binary vectors and 0 being no correlation between binary vectors. The significance of each score was assessed using a permutation test, and scores where *p* < 0.05 were reported.

## Results

In this study we used tools from community ecology to look for spatial structure in the TE communities of 11 genomes. we found that ~ 60% of the variation in TE communities can be explained by spatial patterns. Furthermore, in the human genome 40% of the variation in the TE community was explained by the 3D structure of the genome, and half of that was explained by the chromosomal environment (genomic subcompartment).

### Window Size

The results of this window size analysis are shown in Figure 1. Spatial patterns were significant predictors of TE community composition in 131 of the 149 chromosomes analyzed at all window sizes (*p* < 0.05). Of the 18 other chromosomes, the 16 *Saccharomyces cerevisiae* chromosomes were only significant at the smallest windows size, 14,753bp at the *p* < 0.05 level. The other two chromosomes that were not significant at all window sizes were the X and Y chromosomes in the *D. melanogaster* genome. Both of those chromosomes were significant at the *p* < 0.05 for window sizes below 100,917pb. The X chromosome was not significant at any larger window size, and the Y chromosome was also significant at the 144,808bp window size but not at any window sizes above this size.

Among the TE communities of the chromosomes that had a significant spatial component an average of ~50% of the variation can be explained by spatial patterns alone, with the highest mean R^2^_adj_ of 60% found in *Danio rerio* and the lowest mean R^2^ of 37% found in *Mus musculus* (Figure 1).

**Figure 1:**
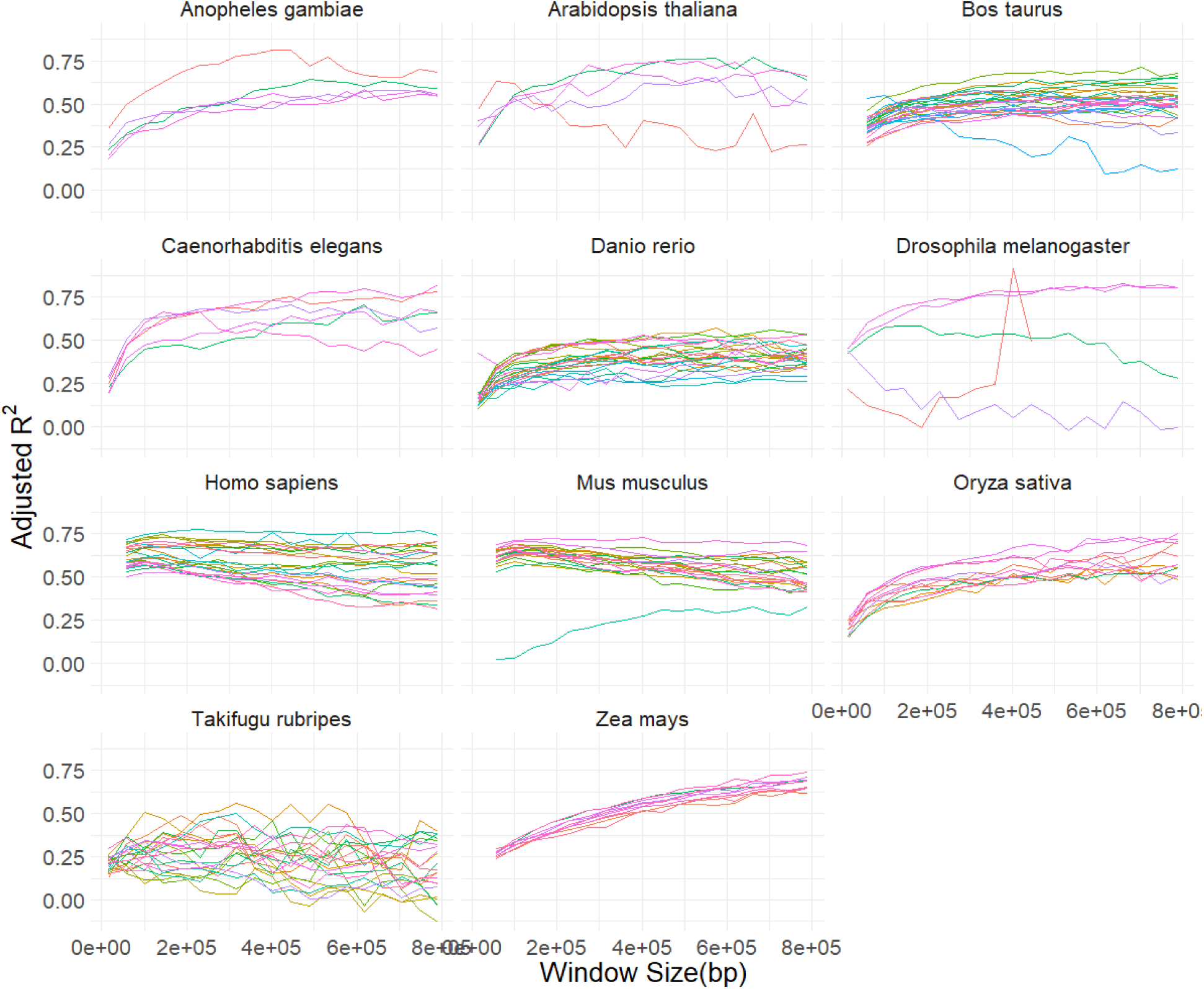
R^2^_adj_ of 11 genomes at multiple window sizes. This figure shows the R^2^_adj_ of for each of the 11 analyzed genomes at each of 20 different window sizes. Each coloured line represents one chromosome and each pane is a different genome. The genomes lacking results for some of the larger window sizes do so because at those sizes the smaller chromosomes in that genome would have been made up of less than three windows.

### Genomic and Chromosomal Properties

The relationship between the average amount of variation in TE communities explained for each genome (R^2^_adj_) and whole genome properties is shown in Figure 2. None of the whole genome properties, Chromosome number, Cvalue, Sequenced length, or the difference between Cvalue and sequence length, showed a significant relationship with average R^2^_adj_ (all *p* values > 0.05).

This remained true after accounting for differences based on phylogeny using phylogenetic independent contrast. After correcting phylogeny with PIC, R^2^_adj_ and GC% were positively correlated (*p* < 0.05), while the other chromosome properties showed no significant correlations with R^2^_adj._

**Figure 2:**
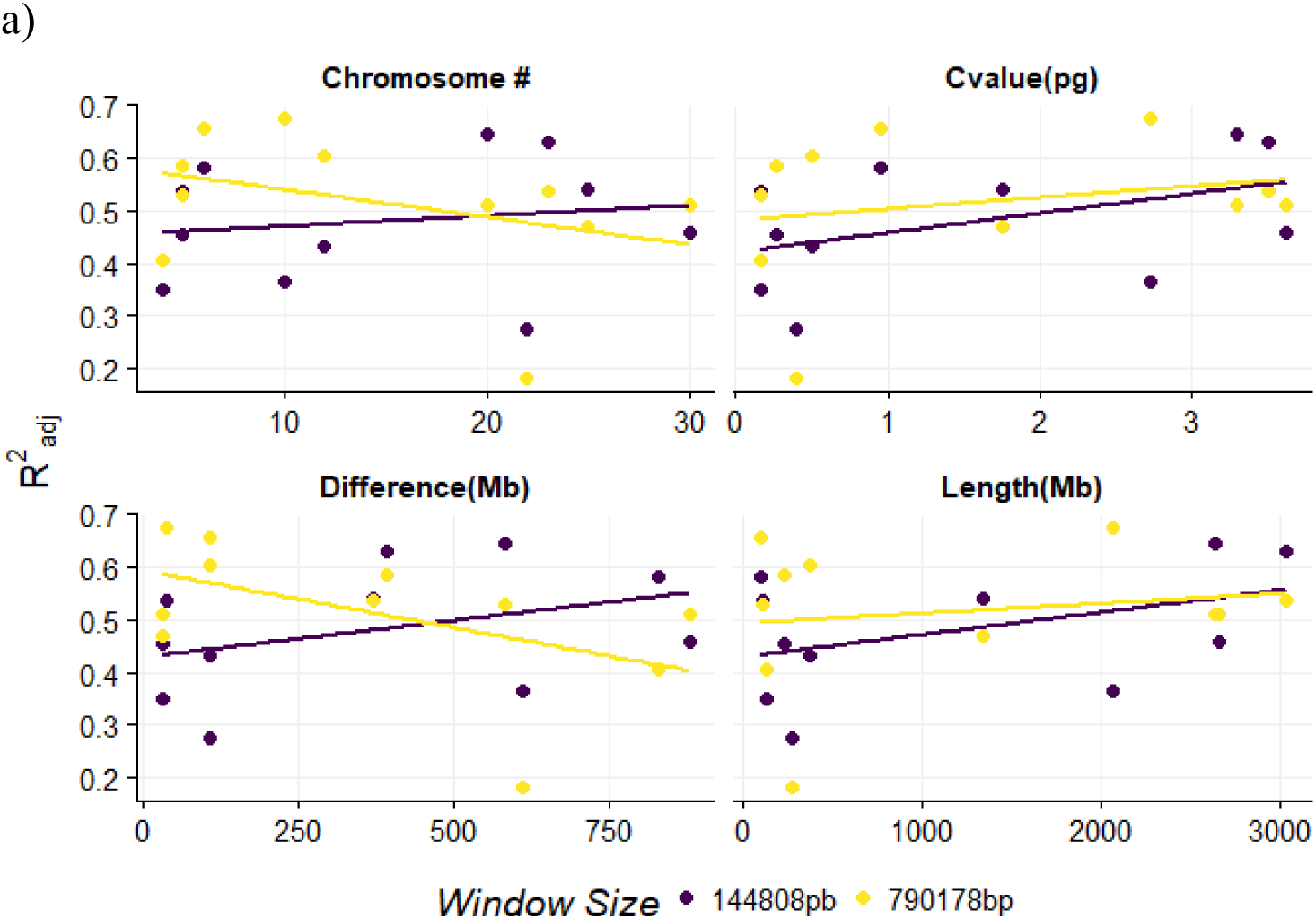
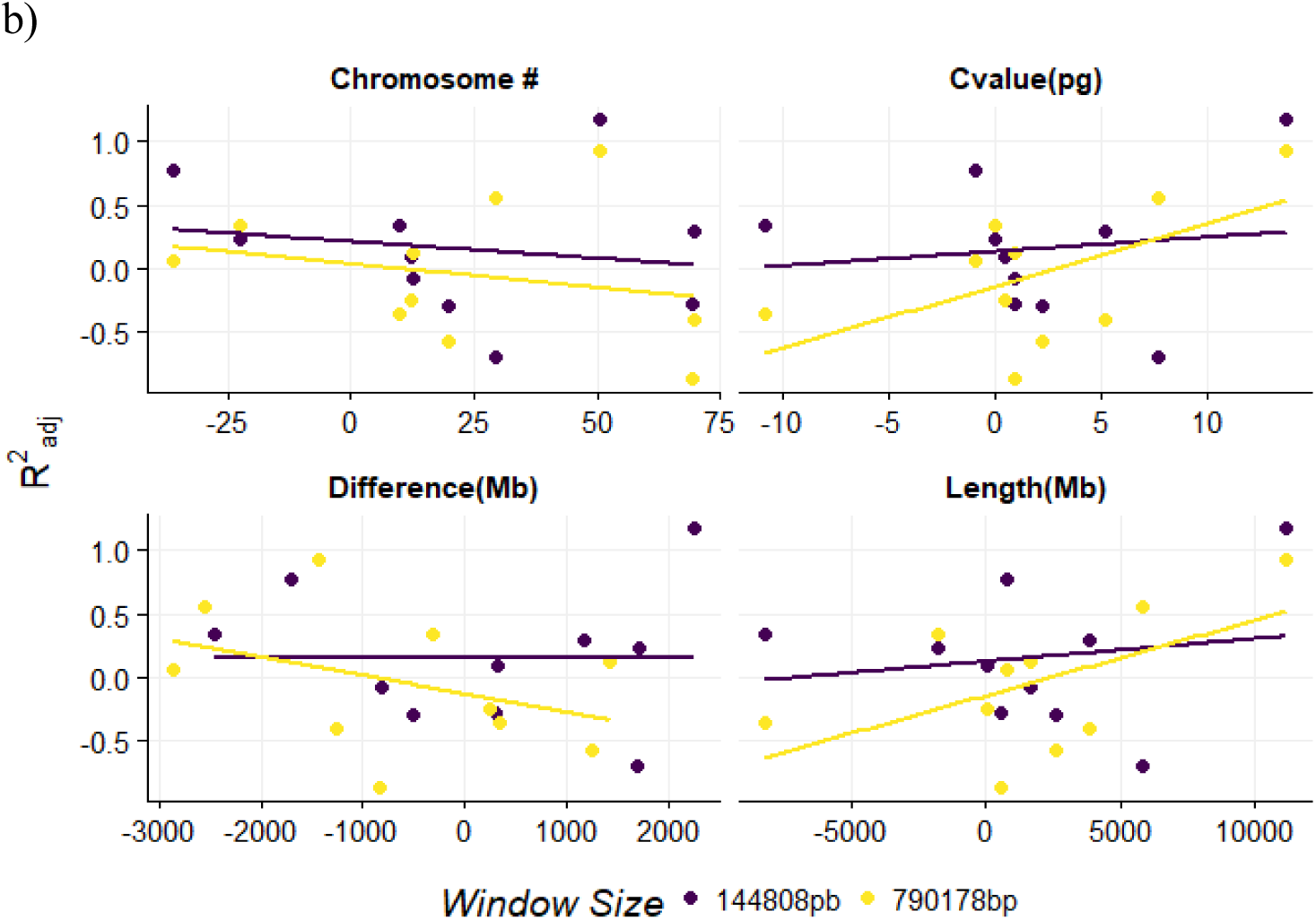
Genome properties versus the R^2^_adj_ across 11 genomes. a) shows the correlation between R2adj and 4 properties of the genome: Chromosome number, Cvalue, sequenced length, and the difference between Cvalue and sequence length. A) Show the raw results, while b) shows the results after PIC analysis. In both cases none of the correlations were significant.

**Figure 3:**
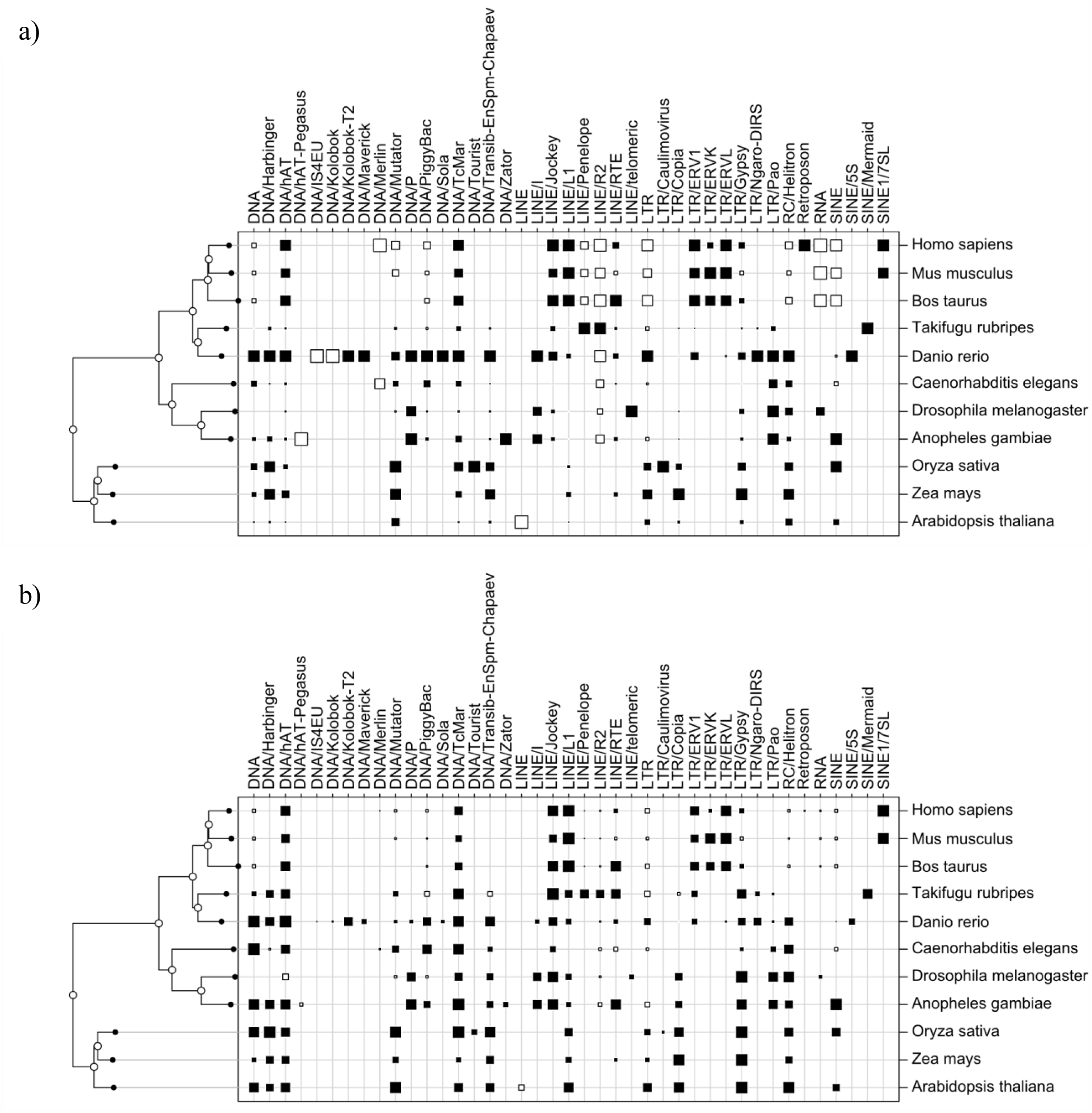
TE presence and abundance in 11 genomes. This figure shows the evolutionary distance between each of the genomes, alongside a table showing which TE superfamilies are present in each genome. The size of each square represents the log of the abundance of each TE family. Black squares represent superfamilies that have at least some of their between-community variation explained by spatial patterns and white squares show superfamilies where no spatial pattern was found. In a) the squares are scaled so that the largest square with each superfamily is the same size. This allows the for the comparisons of superfamilies that have different higher or lower average numbers across genomes. In b) the squares are normalized so that the largest square in each genome are the same size to allow for comparisons across genomes with vastly different numbers of TEs.

### Spatial Importance of 3D Spatial Structure

The 3D spatial structure measured by the HiC interaction frequency explained on average 43%±12% of the TE community distribution within each human chromosome. This was always a subset of the variation explained by our initial analysis in which distances were generated using the PCNM procedure (R^2^_adj_ 69%±7%).

Of the variation explained by HiC data, nearly half of that variation (a total of 22%±12%) is explained by chromosome subcompartments. An additional 7%±4% is explained by the type of subcompartment, but not by HiC data. Gene location data was also analyzed, however it only explained a total 2%±3% of the variation in TE community (Figure 4).

**Figure 4:**
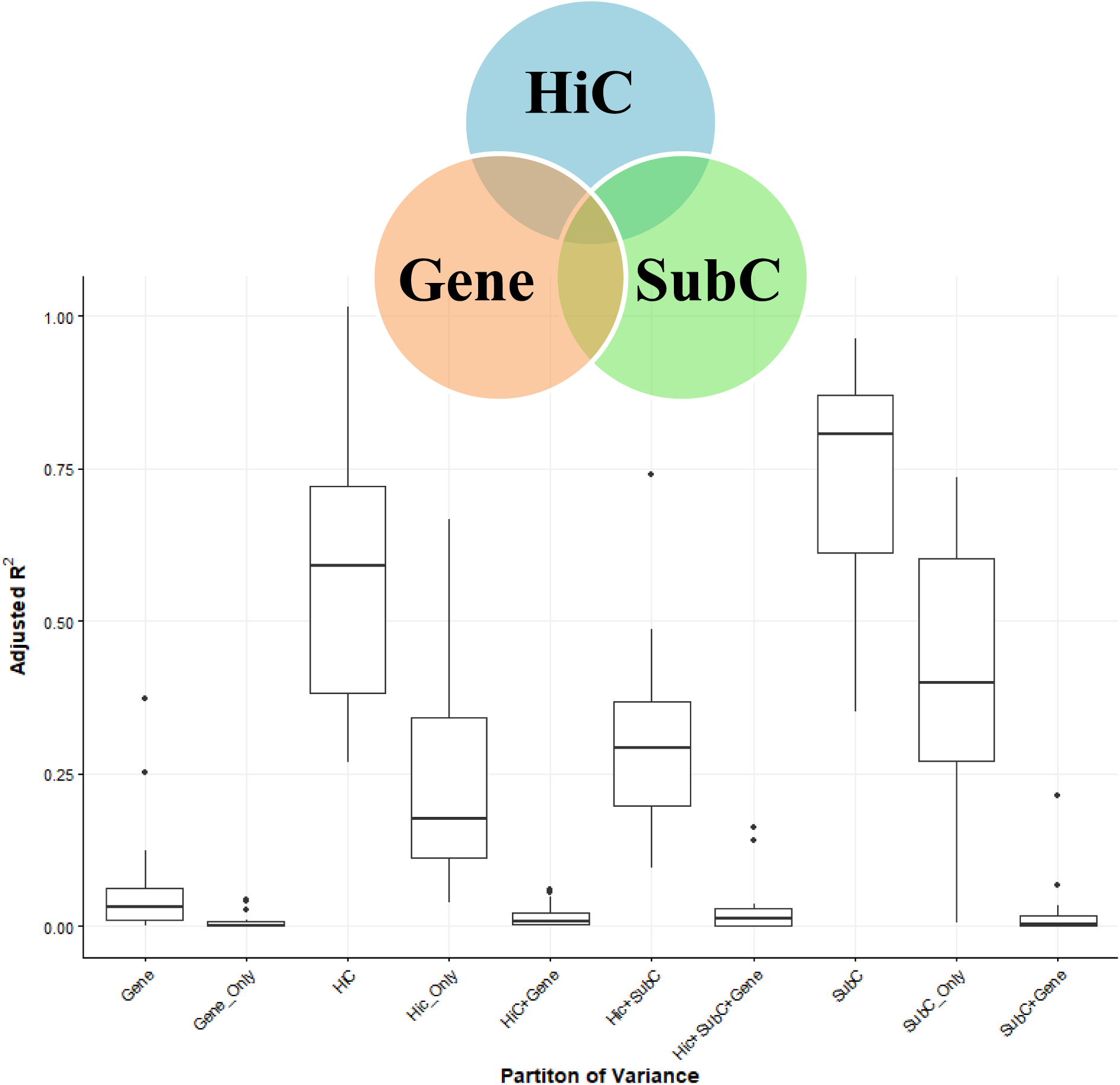
Amount of Variation in TE community composition explained by each environmental factor in the human genome. This figure shows a breakdown of the R^2^_adj_ for the TE communities of each chromosome in the human genome by what environmental factor explains that variation. In this case R^2^_adj_ indicates the amount of variation in the TE community explained by each variable. The R^2^_adj_ for the TE communities of each chromosome are partitioned into those explained by 3 explanatory variables. 1) Number of Genes in each window. 2) Which subcompartments the window was made up of and 3) How close the windows were, as measured by HiC interaction frequency. Each boxplot in the bottom panel represents one of the sections in Venn diagram above the above. The Gene, HiC and SubC boxplots represent the whole circle in the Venn diagram, while the remaining boxplots represent the 7 subsections.

### Indicator Species Analysis

Indicator value (IndVal) scores were produced for each TE within each of 22 human chromosomes. The mean Indval score across all chromes was .55, with the scores ranging from the highest DNA/hAT-Tip100 and LINE/RTE (IndVal= .95 and .89 as indicators for subcompartment B2 or NA on chromosome 22), to the lowest, satellite/centromere (IndVal = .35 for subcompartment NA on chromosome 16) (Figure 5a and Table S1).

Phi scores were also produced for each of the 22 human chromosomes. This resulted in 93 significant potential indicator TEs among the 22 chromosomes. The mean Phi score across all chromes was .38, with the scores ranging from the highest DNA/hAT-blackjack (IndVal= .82 for subcompartment A2 on chromosome 21), to the lowest, rRNA (Phi = .23 for subcompartment B1 on chromosome 14)(Figure 5 b and Table S2).

Overall, the consistency of indicator scores between chromosomes was low, as shown in the lower panels of Figure 5 a and b. The majority of TEs were not significant indicators on more than 1-2 different chromosomes, and those that were indicators on multiple chromosomes were rarely indicators of the same environment type (Figure 5a and b, lower panel). The exceptions to this were found in the Phi scores of Alu and scores for satellite DNA. The Phi scores of Alu showed a significant correlation with an environment on 8 chromosomes, with 4 of those correlations associated with the A1 subcompartment and a fifth being with the A1+NA subcompartment pair. The various categories of satellite DNA showed a more consistent pattern. When the Phi/IndVal score was significant, Satellite DNA was always an indicator of the NA subcompartment. For centromeric satellite DNA, this relationship was detected in 5 chromosomes by the Phi score and 3 chromosomes by the IndVal score.

**Figure 5:**
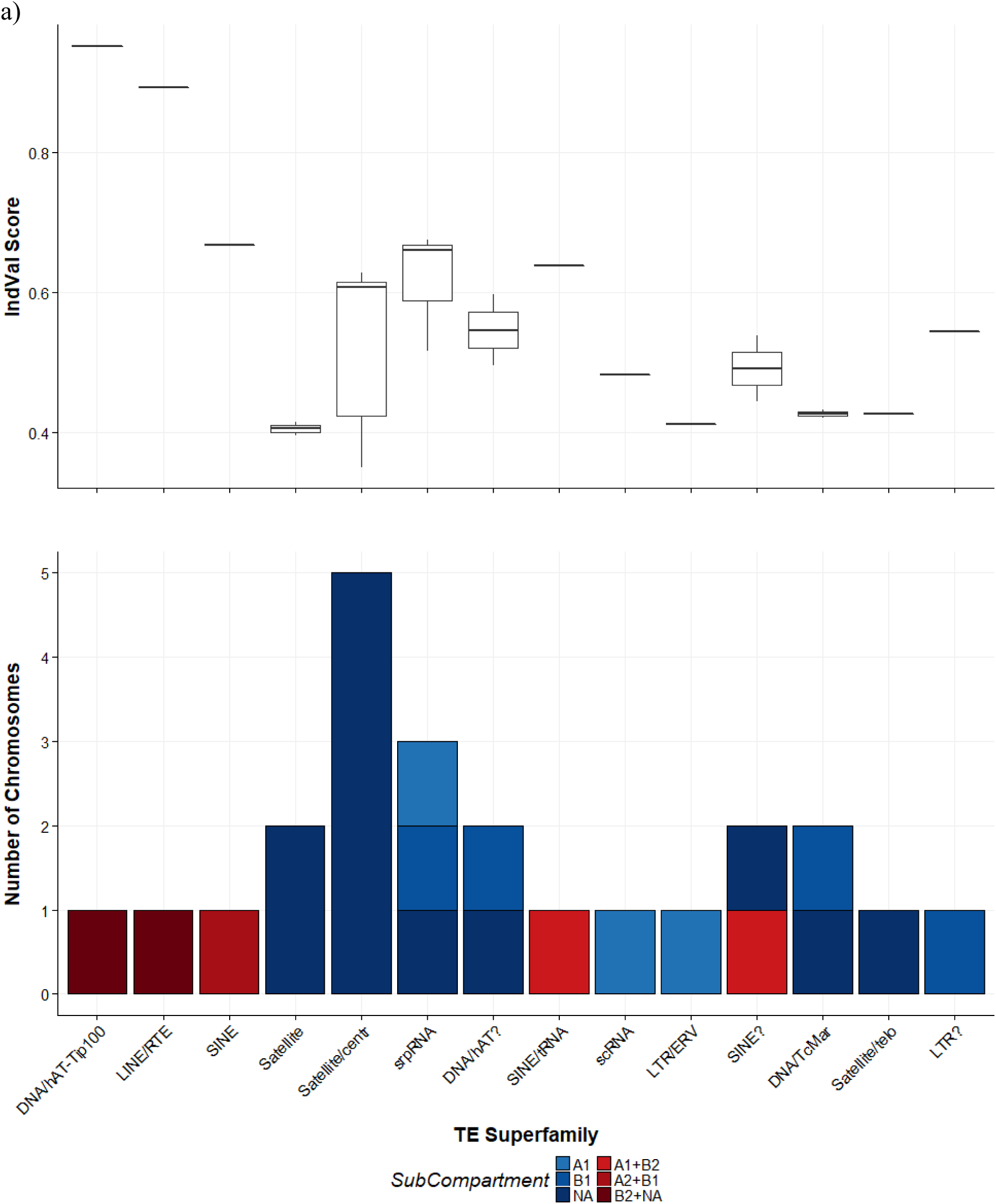
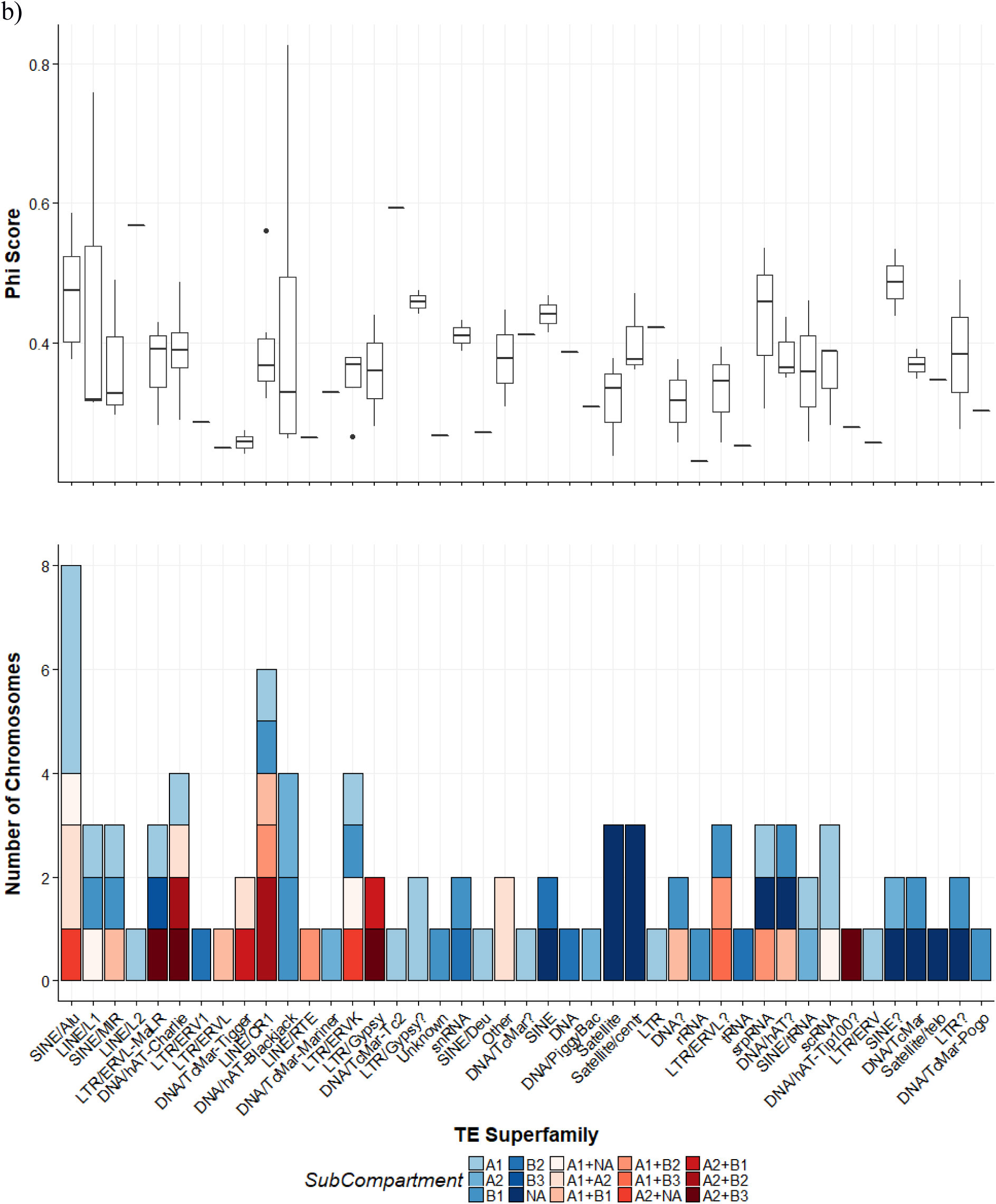
Results of indicator analysis produced by multipatt function from R package vegan. a) Shows the IndVal scores resulting from then analysis. b) Shows the Phi scores resulting from the analysis. For both a) and b) the top boxplot shows the distribution of IndVal/Phi scores, while the lower stacked bar plot shows how often TEs are indicators of a given environment. Blue colors are scores for single environments while reds are environment pairs.

## Conclusions

The underlying spatial structure that present in these linear TE communities, like the underlying spatial structure of communities consisting of organisms, can only be explained by a complex mix of both evolutionary and ecological factors. In TE communities, these patterns are further complicated by selection pressures occurring at both the level of the host and the level of the TE, which can work to either reinforce or counteract each other. This complexity necessitates careful consideration of both evolutionary and ecological processes, and the scale at which they are acting, before making conclusions about TE communities. At the host level, ecology focuses on interaction between different species, which rarely if ever involves TEs. Evolutionary processes at the host level can involve TEs, but mainly as sources of mutation, as they cause changes in the focal entity, the host, or by host level processes affecting TEs, such as drift fixing TE insertions in small populations. TE evolutionary processes are those in which the TEs themselves are changing. This is the focus of most TE research. This analysis focused on the ecology at the level of the TE, by examining the relationships between various types of TEs and their environment. Although TE ecology is often discussed, the boundaries between these levels and processes are often not considered before designing experiments or making conclusions, violating many of the assumptions for those processes [32,61].

The analysis presented by Saylor *et al.* [33] focused on explicitly ecological methods adapted from community ecology. That study demonstrated the utility of such an approach in principle and in practice. In this paper, we continued that analysis by extending this proof-of-concept in four major ways: 1) By examining the impact of window size selection in the implementation of the method; 2) by applying the approach to 11 genomes of varying sizes and compositions; 3) by considering additional genomic and chromosomal factors that may influence TE abundance and distribution; and 4) by examining the role of how physically close areas of the genome are in predicting TE community.

### Consistency of Results Across Genomes and Window Sizes

Importantly, the results obtained using various window sizes and multiple genomes were remarkably consistent, suggesting that this approach will be broadly applicable in analyzing TE abundances and distributions across a wide range of taxa. In particular, the present analyses demonstrated that a large amount of the spatial variation in the TE community of each genome was explained by accounting for the spatial distribution location of those TEs. In other words, a large proportion of the TEs likely to be found in a section (window) of any chromosome can be predicted based on the relative location of that window along the chromosome.

Care must be taken when adapting any set of tools for a new use. The analysis of TE communities using methods from community ecology appears promising, as the spatial location of TEs was correlated to the composition of the TE community in all 193 chromosomes analyzed across all 11 genomes. Although the spatial distribution of TEs was significant on each chromosome, the amount of the TE community in each window that could be explained in this way was not, and the TE superfamilies that were spatially structured were not consistent across chromosomes. One would expect that decreasing the window size would increase the explanatory power of spatial patterns as it would allow finer-scale spatial patterns to be detected. However, reducing the window size too much results in a steep drop-off in explanatory power, which is most evident in the smaller genomes (Figure 1). This fact highlights one notable limitation of the method when applied to genomes vs. ecological communities. Ecological communities typically have less complete sampling, and the statistical methods used on this ecology data are designed with this in mind. In the context of analyzing whole genome sequences, there is a risk of creating statistical overpower as the degrees of freedom are extremely high, which can cause the model to falsely identify patterns as significant.

This was a known issue in the original ecological application of the PCNM method, and the solution was to use R^2^_adj_ instead of the raw R^2^ value (Equation 1). This R^2^_adj_ value reduces the R^2^ based on the inflated statistical power; however, the hundreds of thousands of observations typical of a whole genome analysis are too extreme for even the R^2^_adj_ calculation, and R^2^_adj_ plummets as the difference between the number of windows and the number of potential spatial patterns generated from the PCNM procedure increases (Figure 6). This brings up an importation point raised by Linquist *et al.* [32], namely that the assumptions and limits of any model need to be carefully considered before being applied to a different type of data. In this case without considering the assumptions and function of the model, one might assume that all TE interactions happen at a very large scale, as R^2^_adj_ is lower in analyses with small window sizes. This may be true in some cases, fine scale patterns in TE distribution may also be obscured by lack of statistical power.

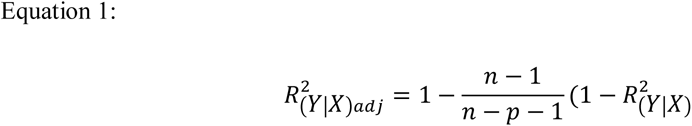

Where n is # of observations (windows) and p is # parameters (Potential PCNMs graphs)

**Figure 6:**
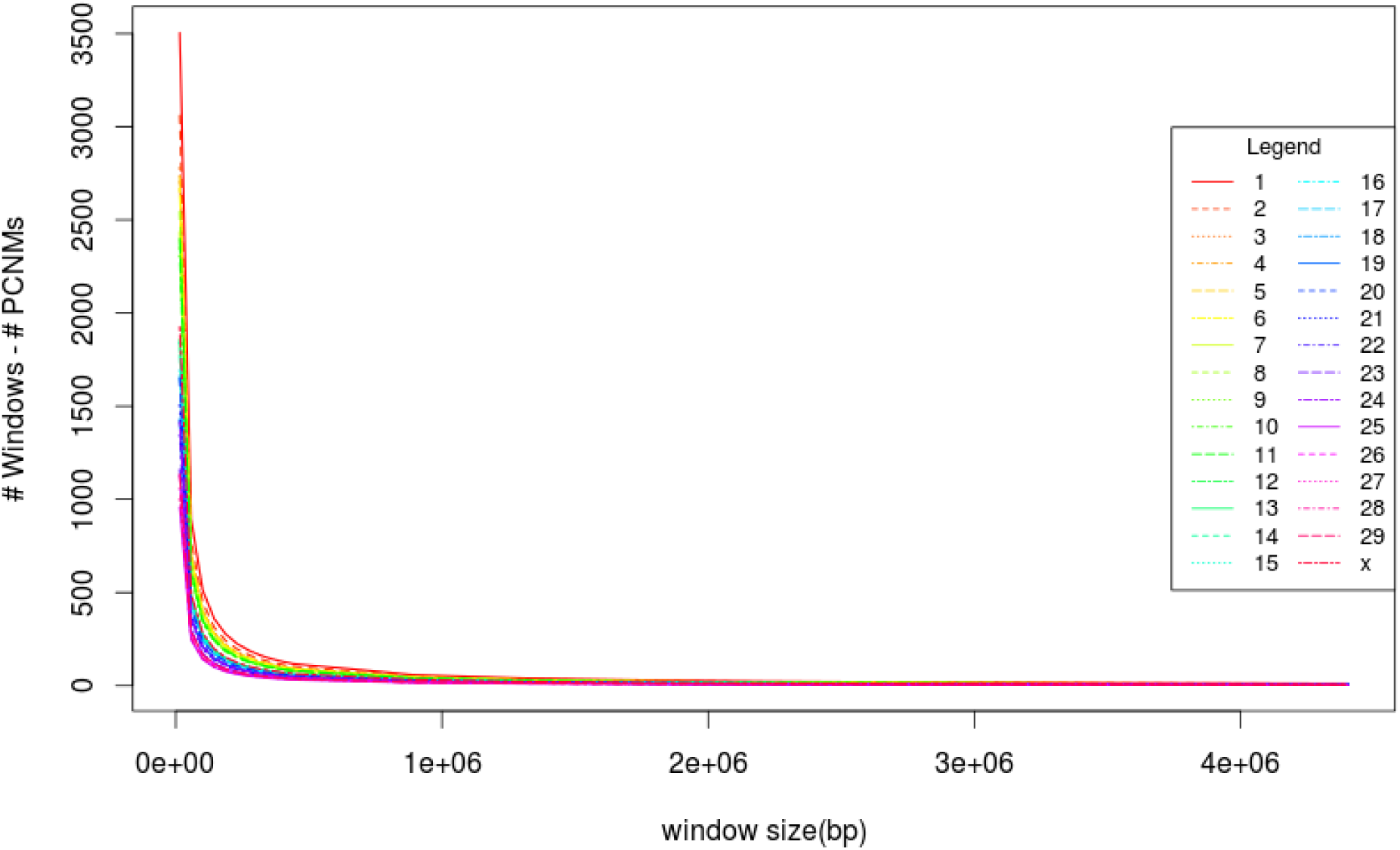
Adjusted degrees of freedom verses window size in RDAs of the *Bos taurus* genome

Despite this limitation, the analysis of 11 genomes differing in size and chromosome number revealed some interesting patterns. Notably, similar TE families were implicated in accounting for spatial variation of the TE community in each individual genome, regardless of the window size used. Moreover, the only TE families were significant at larger window sizes and became non-significant at smaller windows sizes appeared to be eliminated due to the larger adjustment to the R^2^_adj_ value. Thus, results of the spatial analysis are relatively robust to window size even across very different genome sizes or numbers of chromosomes. By the time window size becomes sufficiently small to engender computational limitations, the majority of the TE families identified as spatially relevant continue to be identified as such on each chromosome. By contrast, those TEs that are not considered significant in terms of spatial structure at certain window sizes are typically the ones that had the lowest explanatory power initially. This robustness notwithstanding, it would still be advisable to implement the analysis multiple times with different window sizes when dealing with previously unstudied genomes as a matter of best practice.

### Patterns of Transposable Element Distribution

The results of the present analyses indicated that spatial patterns explain ~50% of the variation between TE communities in each of 11 distantly related genomes, and that larger chromosomes exhibit more spatially structured TE communities. This consistency suggests that there are some common factors influencing the locations of TEs within a given genome. The cause(s) of the observed spatial patterns is still not completely clear, however our evidence suggests that the genomic environment itself may play some role. This is shown by TEs with similar abundances in different genomes being spatially structured in one genome, but not in another (Figure 3a). These differences in amount of spatial structure in different genomes may indicate that the same TEs in different genomes may be found in different patterns of differing strengths based on the different environment – in this case the genome. Although a genome’s properties were not related to the amount of spatial organization of that genome’s TE community, it was related to the composition of the TE community. This is shown in the difference between the number and identity of the TE superfamilies organized by a spatial pattern in different genomes. For example, organisms that are closely related phylogenetically have similar groups of TEs that are spatially structured. The TE superfamilies found in plants were almost all spatially structured, while in mammals only about half of the superfamilies were shown to have some degree of spatial structure (Figure 3).

This large-scale difference among taxa could be explained in several different ways, including: 1) a shared history of TE insertions among similar closely related genomes, 2) interactions between TEs and their genomic environment, which are more similar in more closely related species; or 3) some properties of the TEs themselves, with the types of elements differing among taxa. It seems likely that options 1 and 2 are connected, as more closely related species are more likely to have more similar genomic environments than more distantly related species. For example, plants and animals have similar but distinct systems to suppress genome activity [62]. As a result of these differences, plants and animals have unique patterns in methylation. In plants, methylation is preferentially found in repetitive areas of the genomes [63], including TEs, the methylation occurs on cytosines irrespective of the sequence surrounding them [64]. In mammals, methylation is found primarily on CG dinucleotides, and rarely in any other context. This methylation is found throughout the genome and is estimated to be found on ~70-80% of mammalian CG sites [65]. These differences could affect how and when individual TEs are suppressed, potentially contributing to particular spatial distributions. TE distribution can also be caused by TE-specific properties, such as insertion site preference. These preferences range from TEs that are enriched in specific regions, to those with very specific target sites. TEs that display insertion preference for specific regions include MITEs in genic regions [24,66], Ty5 elements in heterochromatic regions of *Saccharomyces* [67,68],Ty3/gypsy elements in the centromeres of plants [69], and the non-LTR elements that maintain chromosome ends in *Drosophila* [3,70,71]. TEs with very specific target sites are often found in various RNA genes, such as Pokey and R2 elements 28s RNA genes, and Dada DNA element family, some of which target U6 and U1 snRNA, and various tRNA genes [72]. In that regard, spatial structure in TE distributions could reflect the spatial patterns of different insertion sites in different genomes.

### 3D Analysis and Indicator Species Analysis

By incorporating frequency of interaction data from high res HiC data we were able to explain 2/3 of the variation in TE community explained by our more complete PCNM model. The variation explained by the HiC data is a subset of the PCNM which generates artificial community abundances in such a way that any spatial pattern between the windows along a chromosome is accounted for. The HiC analysis is a specific subset of these based on the frequencies at which the windows along the chromosome interact. The HiC dataset explaining 2/3 of the variation means that a large part of the variation in the TE community is explained by the physical closeness of the sections of the chromosome when they are uncondensed in the nucleus. Additionally, half of the community data explained by the HiC data is also explained by genomic subcompartments. Areas of the chromosome that have been classified as the same compartments have been shown to have consistent epigenetic marks, which play a role in how accessible these areas are to specific TEs [68,73,74]. Based on the banding patterns of the HiC analysis, they also form clusters of chromosome loops, the borders of which are physically bound together with CTCF anchor protein (Figure 7).

**Figure 7:**
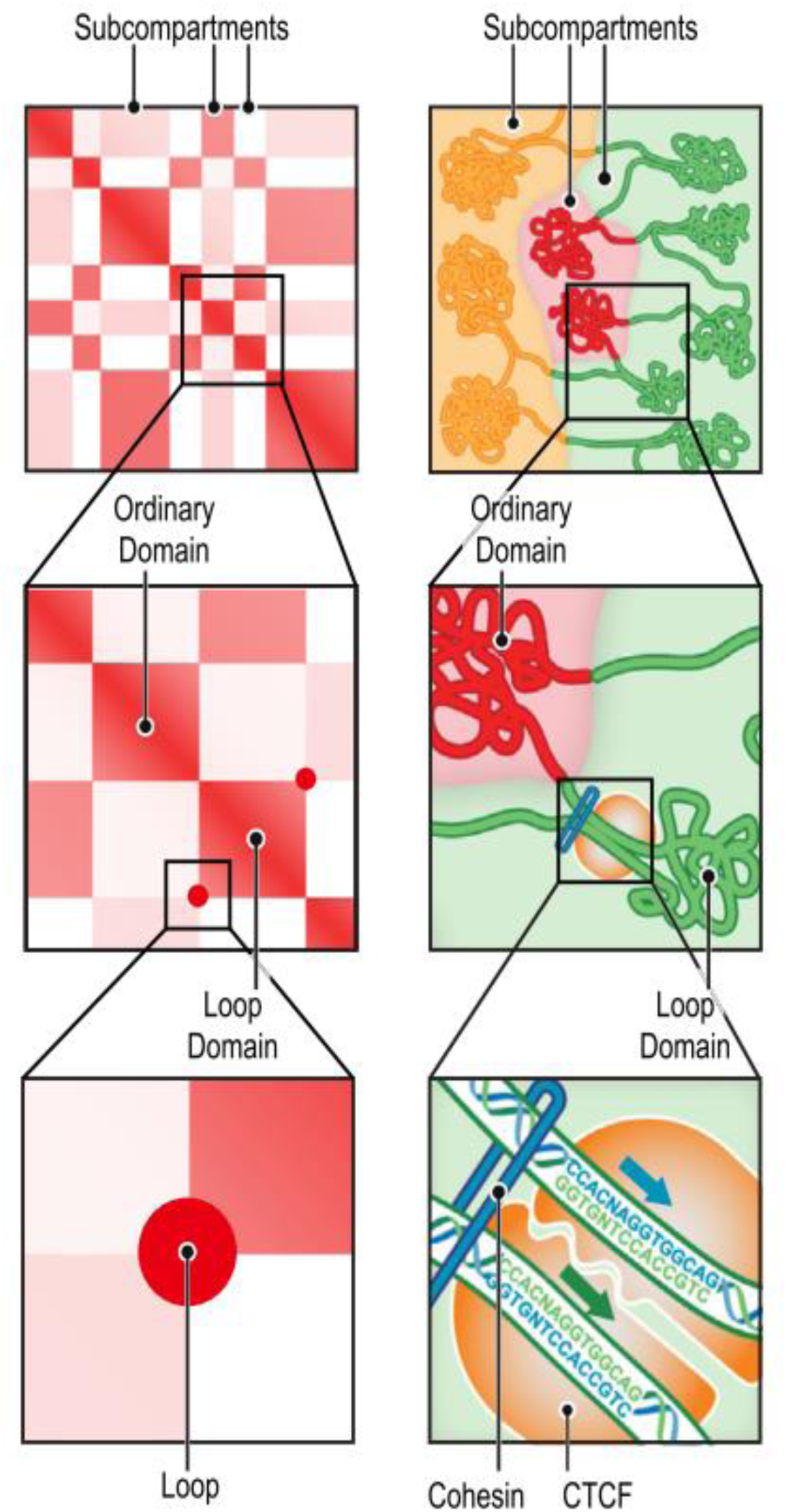
Subcompartment diagram. Shows how subcompartments are made up of chromosome loops, the boundaries of which are bound together with CTCF anchor proteins. Reproduced from [40] Figure 1d.

Knowing that genomic subcompartments were able to explain a large amount of the variation in TE communities, we examined the predictive power of this environmental variable on the TE community. Figure 5 shows far fewer significant IndVal scores than Phi scores, and that neither of these scores were as high as one would want to see for use as a traditional indicator used in something like biomonitoring. The differences in the number of significant scores is likely that the TEs tend not to be found exclusively in one environment. The IndVal score weights this specificity more heavily than the Phi score, which is a measure of correlation [60]. Thus, our results show TE superfamilies that are found in greater abundance in some subcompartments than others. For example, Figure 5b shows that Alu elements are found most often in A1 or A1+NA subcompartments on five different chromosomes.

Overall, with the exception of Alu, and the NA compartment, which seems to be associated with satellite DNA, the TE superfamilies identified by the IndVal and Phi analysis were not consistent across chromosomes (Figure 5). This inconsistency indicates that the ecological patterns structuring TE communities does not extend to the chromosome level. This indicates that transposon ecology may need to think of TE communities at a more local scale than that of the genome, which is currently the standard (e.g. see[75–78]). Instead the TE communities may be structured at a smaller scale, with the TEs of a whole chromosome, or a whole genome, being more analogous to a metacommunity, with dispersal occurring between more local communities, or from the metacommunity (the TE population of the chromosome/genome) at large [79].

### Final Remarks and Future Directions

By taking an ecological approach to TEs and drawing inspiration from existing ecological methods, we found that the distribution of ~50% of the TEs, within a diverse set of genomes, is distributed along the chromosome in a detectable pattern. Across these genomes, the TE superfamilies in which these patterns are detectable are correlated with the phylogeny of the host taxa. In a more focused analysis of the impact of 3D spatial relationships on TE community, we found that a large part of TE community composition was structured by physical distance between the communities, and the genomic subcompartment the community was found in. From those results, we suggest that, along with producing and examining more high resolution genomic HiC data, in order to more explicitly define the scale of TE communities, those interested in the ecology of the genome should continue to look at community ecology, and perhaps more specifically metacommunity theory, to better understand the distribution of TEs within and between genomes.

## Declarations

### Ethics approval and consent to participate

Not applicable

### Consent for publication

Not applicable

### Availability of data and material

The reference genome sequences use in this analysis are available on the GenBank website under the DOIs found in Table 1

The HiC interaction produced by Rao *et al.* [40] is available from the Gene Expression Omnibus (GEO) database GEO accession GSE63525

The code use to generate the results will be posted on is being contributed to GitHub. A link will be provided by the time the paper is published by the time of submission

### Competing interests

The authors declare that they have no competing interests

### Funding

Supported by Natural Sciences and Engineering Research Council of Canada (NSERC) Discovery grants to SCK, TRG, and KC.

### Authors’ contributions

BS designed analysis with input from CK, TRG, and SCK. BS implemented the analysis, wrote code, and the manuscript. CK, TRG, and SCK were all involved in editing manuscript.

## Acknowledgements

We would to thank the genome ecology working group for development of the genome ecology framework this analysis was inspired by, and the Compute Canada high performance computing cluster for the resources required to run the analysis.

## Supplementary Tables

**Table S1:** Significant IndVal scores for each TE on each produced by the indicator species analysis

| *Chromosome* | *IndVal* | *p value* | *TE* | *SubCompartment* |
| --- | --- | --- | --- | --- |
| 1 | 0.421272 | 0.047 | DNA/TcMar | NA |
| 1 | 0.538299 | 0.023 | SINE? | NA |
| 2 | 0.544331 | 0.001 | LTR? | B1 |
| 3 | 0.412272 | 0.024 | LTR/ERV | A1 |
| 3 | 0.444923 | 0.042 | SINE? | A1+B2 |
| 3 | 0.660182 | 0.004 | srpRNA | A1 |
| 4 | 0.59728 | 0.002 | DNA/hAT? | NA |
| 6 | 0.41502 | 0.005 | Satellite | NA |
| 7 | 0.396417 | 0.015 | Satellite | NA |
| 7 | 0.628799 | 0.001 | Satellite/centr | NA |
| 8 | 0.638628 | 0.005 | SINE/tRNA | A1+B2 |
| 11 | 0.607026 | 0.002 | Satellite/centr | NA |
| 12 | 0.615626 | 0.026 | Satellite/centr | NA |
| 13 | 0.482719 | 0.041 | scRNA | A1 |
| 14 | 0.433492 | 0.004 | DNA/TcMar | B1 |
| 16 | 0.350686 | 0.026 | Satellite/centr | NA |
| 18 | 0.675529 | 0.007 | srpRNA | NA |
| 19 | 0.423761 | 0.004 | Satellite/centr | NA |
| 19 | 0.426401 | 0.006 | Satellite/telo | NA |
| 20 | 0.667816 | 0.026 | SINE | A2+B1 |
| 20 | 0.515964 | 0.034 | srpRNA | B1 |
| 21 | 0.951528 | 0.013 | DNA/hAT-Tip100 | B2+NA |
| 21 | 0.892915 | 0.037 | LINE/RTE | B2+NA |
| 22 | 0.495763 | 0.039 | DNA/hAT? | B1 |

**Table S2:**
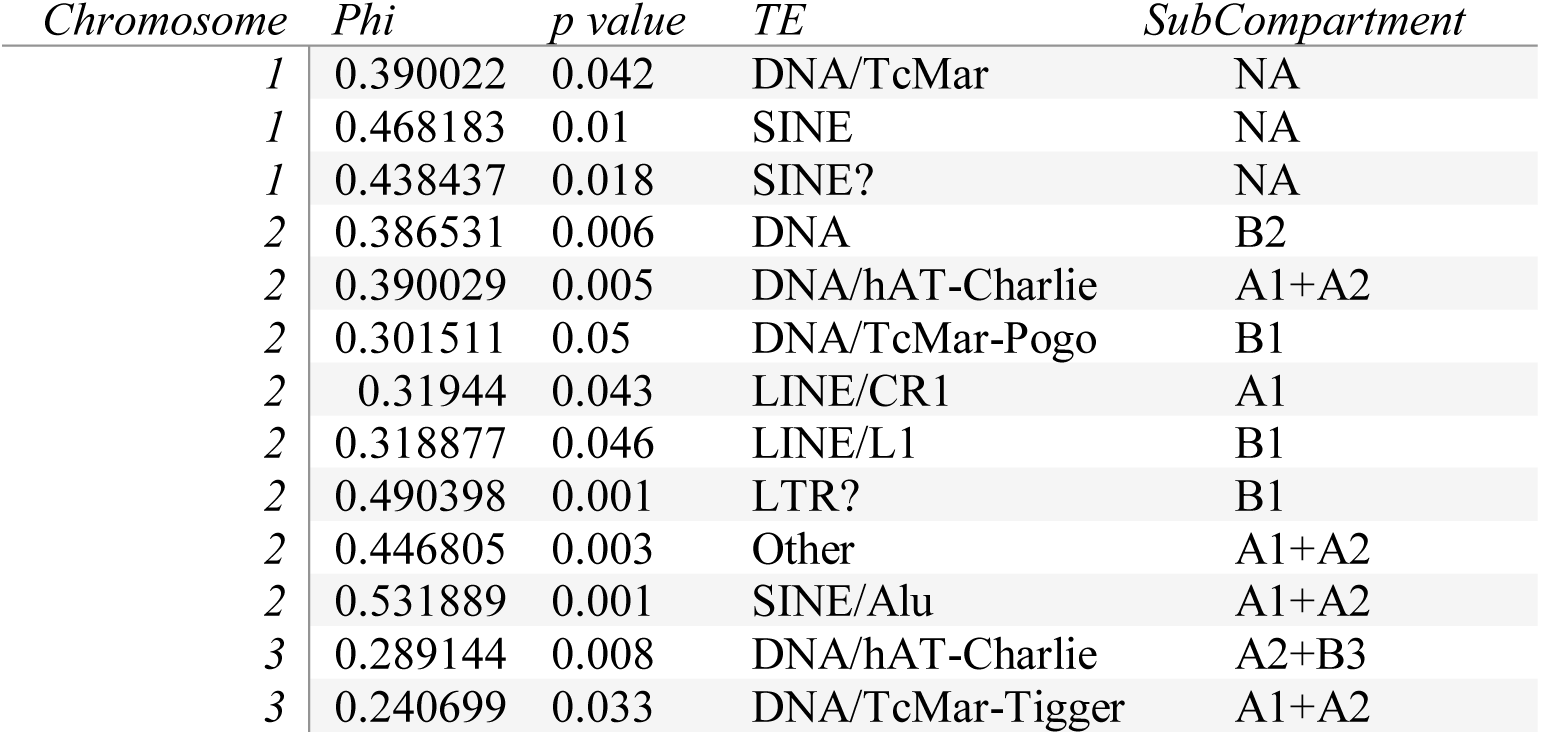
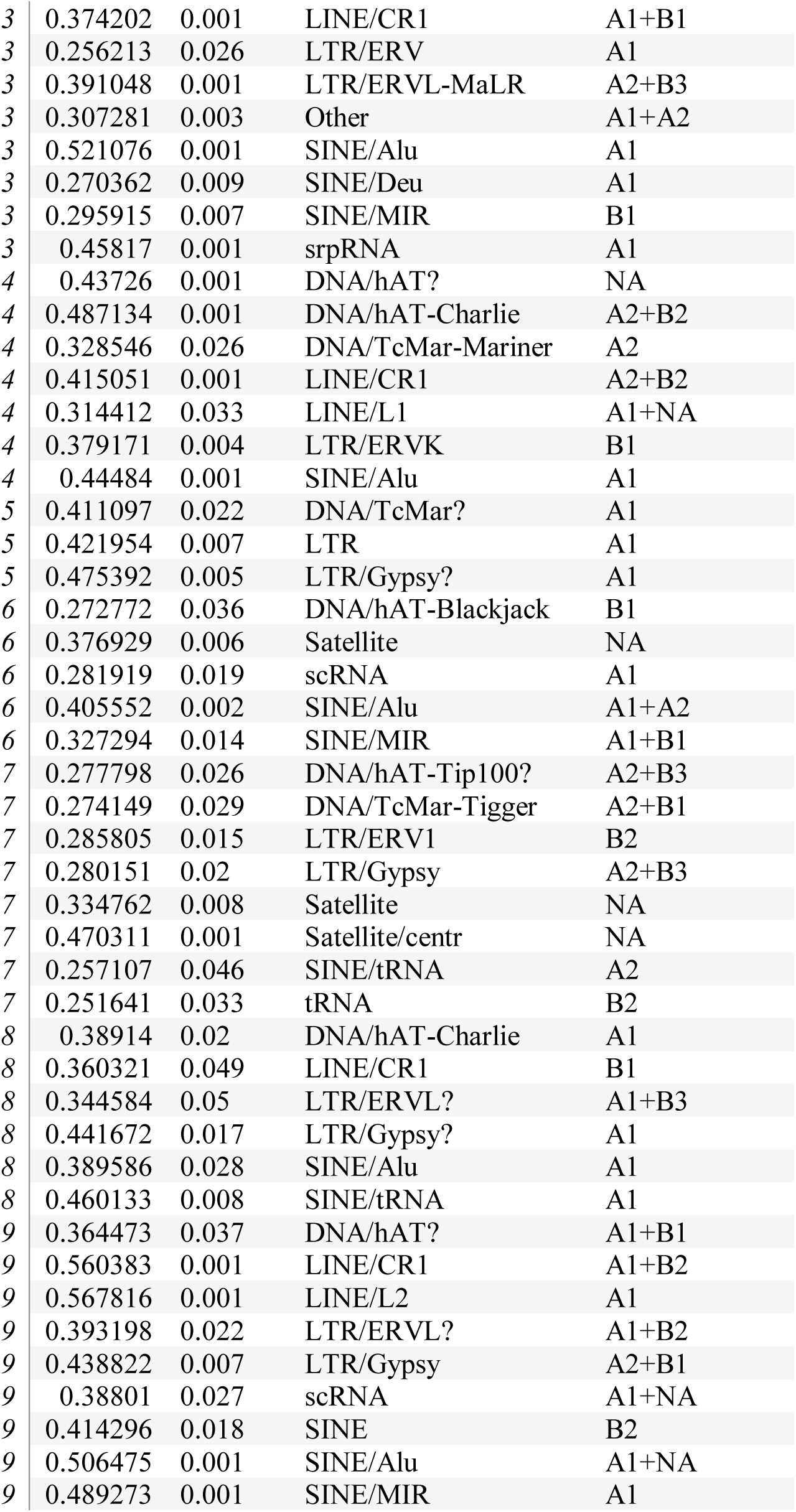
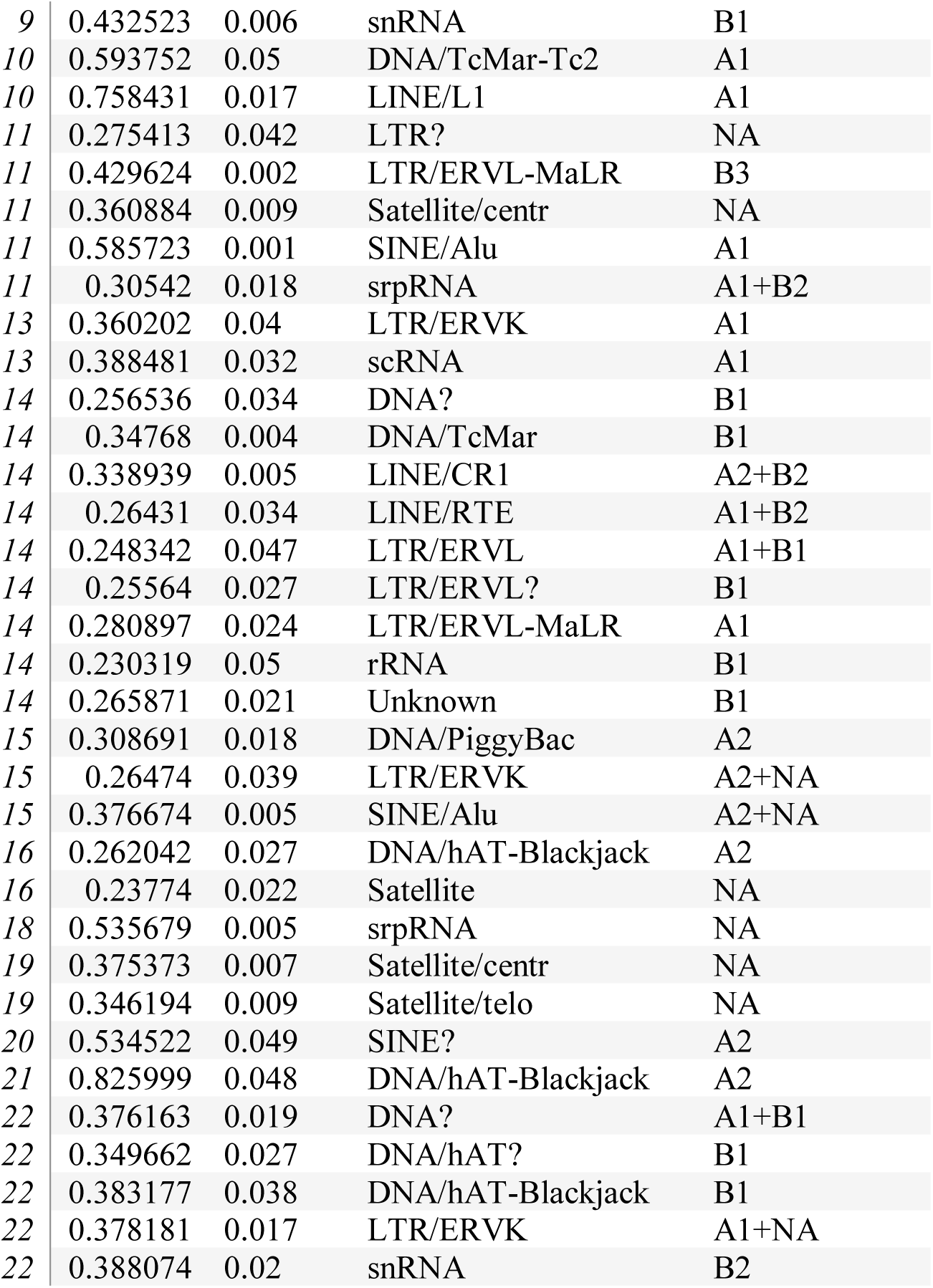
Significant Phi scores for each TE on each produced by the indicator species analysis

